# Activation of RIG-I-mediated antiviral signaling triggers autophagy through the MAVS-TRAF6-Beclin-1 signaling axis

**DOI:** 10.1101/340323

**Authors:** Na-Rae Lee, Junsu Ban, Noh-Jin Lee, Chae-Min Yi, Jiyoon Choi, Hyunbin Kim, Jong-Kil Lee, Jihye Seong, Nam-Hyuk Cho, Jae U. Jung, Kyung-Soo Inn

**Affiliations:** Department of Fundamental Pharmaceutical Sciences, Graduate School, Kyung Hee University, Seoul, Republic of Korea; KHU-KIST Department of Converging Science and Technology, Graduate School, Kyung Hee University, Seoul, Republic of Korea; Convergence Research Center for Diagnosis, Treatment and Care System of Dementia, Korea Institute of Science and Technology (KIST), Seoul, Republic of Korea; Department of Biomedical Sciences, Seoul National University College of Medicine, Seoul, Republic of Korea; Department of Microbiology and Immunology, Seoul National University College of Medicine, Seoul, Republic of Korea; Department of Molecular Microbiology and Immunology, Keck School of Medicine, University of Southern California, Los Angeles, California, United States of America

**Author notes:** Address correspondence to Kyung-Soo Inn.

## Abstract

Autophagy has been implicated in innate immune responses against various intracellular pathogens. Recent studies have reported that autophagy can be triggered by pathogen recognizing sensors, including Toll-like receptors and cyclic guanosine monophosphate-adenosine monophosphate synthase, to participate in innate immunity. In the present study, we examined whether the RIG-I signaling pathway, which detects viral infections by recognizing viral RNA, triggers the autophagic process. The introduction of polyI:C into the cytoplasm, or Sendai virus infection, significantly induced autophagy in normal cells but not in RIG-I-deficient cells. PolyI:C transfection or Sendai virus infection induced autophagy in the cells lacking type-I interferon signaling. This demonstrated that the effect was not due to interferon signaling. RIG-I-mediated autophagy diminished by the deficiency of mitochondrial antiviral signaling protein (MAVS) or tumor necrosis factor receptor-associated factor (TRAF)6, showing that the RIG-I-MAVS-TRAF6 signaling axis was critical for RIG-I-mediated autophagy. We also found that Beclin-1 was translocated to the mitochondria, and it interacted with TRAF6 upon RIG-I activation. Furthermore, Beclin-1 underwent K63-polyubiquitination upon RIG-I activation, and the ubiquitination decreased in TRAF6-deficient cells. This suggests that the RIG-I-MAVS-TRAF6 axis induced K63-linked polyubiquitination of Beclin-1, which has been implicated in triggering autophagy. Collectively, the results of this study show that the recognition of viral infection by RIG-I is capable of inducing autophagy to control viral replication. As deficient autophagy increases the type-I interferon response, the induction of autophagy by the RIG-I pathway might also contribute to preventing an excessive interferon response as a negative-feedback mechanism.

**Importance:** Mammalian cells utilize various innate immune sensors to detect pathogens. Among those sensors, RIG-I recognizes viral RNA to detect intracellular viral replication. Although cells experience diverse physiological changes upon viral infection, studies to understand the role of RIG-I signaling have focused on the induction of type-I interferon. Autophagy is a process that sequesters cytosolic regions and degrades the contents to maintain cellular homeostasis. Autophagy participates in the immune system, and has been known to be triggered by some innate immune sensors, such as TLR4 and cGAS. We demonstrated that autophagy can be triggered by the activation of RIG-I. In addition, we also proved that MAVS-TRAF6 downstream signaling is crucial for the process. Beclin-1, a key molecule in autophagy, is translocated to mitochondria, where it undergoes K63-ubiquitination in a TRAF6-dependent manner upon RIG-I activation. As autophagy negatively regulates RIG-I-mediated signaling, the RIG-I-mediated activation of autophagy may function as a negative-feedback mechanism.

## Introduction

Autophagy is a process that sequesters cytosolic regions and delivers their contents to the lysosomes for subsequent degradation. Both extracellular stimuli, such as starvation and hypoxia, and intracellular stresses, including the accumulation of damaged organelles, induce autophagy to degrade long-lived proteins and damaged organelles in order to recycle and maintain cell homeostasis (1). As autophagy is triggered by infection with intracellular pathogens, such as bacteria and viruses, it is recognized as part of the innate immune system to control and eliminate infections by engulfing and degrading intracellular pathogens (i.e., xenophagy) (2–4). Recent extensive studies have revealed that several autophagic adaptors, such as sequestosome 1 (SQSTM1/p62), optineurin, and nuclear dot protein 52 kDa, specifically recognize the intracellular presence of bacteria, including *Salmonella, Shigella, Listeria*, and *Mycobacteria*, and induce autophagy (5–8). Adaptor proteins, referred to as sequestosome 1/p62-like receptors (SLRs), directly recognize ubiquitinated microbes as their targets to induce autophagy. They are now regarded as a class of pattern recognition receptors of the innate immune system (9).

Besides the SLRs, the innate immune system utilizes a limited number of sensors including Toll-like receptors (TLRs), Nod-like receptors, cyclic guanosine monophosphate-adenosine monophosphate synthase (cGAS), and RIG-I-like receptors (RLRs) to detect various pathogen-associated molecular patterns (10–14). Among these sensors, RIG-I and MDA5 recognize viral RNAs to mount an antiviral immune response. Upon recognition, RIG-I and MDA5 translocate to mitochondria to interact with the mitochondrial antiviral signaling protein (MAVS)/IPS-1/Cardif, a downstream mitochondrial signaling protein. The subsequent recruitment of signaling molecules, including tumor necrosis factor receptor-associated factor (TRAF)3 and TRAF6, results in the activation of transcription factors, such as IRF3/7, NF-κB, and AP-1, leading to the production of type-I interferon.

As autophagy can be activated by infection with pathogens, it is conceivable that the innate immune sensors regulate the autophagic process upon recognition of pathogens. Indeed, the activation of innate immune signaling pathways triggered by innate immune sensors, including TLR4 and cGAS, activates autophagy, indicating that the innate immune system modulates autophagy directly (15–17). Besides participating in the innate immune system by directly eliminating pathogens, autophagy also plays crucial roles in regulating this system to prevent excessive responses (18–20).

A recent study showed that the absence of autophagy amplified RIG-I signaling due to increased mitochondrial MAVS and reactive oxygen species from damaged mitochondria, implicating autophagy in RLR signaling (21). However, it has not been reported whether RIG-I-mediated antiviral signaling directly regulates autophagy. Herein, we show that the activation of RIG-I by its ligands provokes autophagy in a downstream MAVS-TRAF6 signaling axis-dependent manner.

## Results

### Activation of the RIG-I signaling pathway activates autophagy

To investigate whether the recognition of viral RNA by RIG-I can trigger autophagy, HEK293T cells were transfected with the RIG-I agonist, polyI:C, a synthetic doublestranded RNA analogue, or infected with Sendai virus (SeV). Both polyI:C transfection and SeV infection increased the level of LC3-II, a lipidated form of LC3 (Fig. 1A). In both the experimental settings, the change was evident 2 h after stimulation (Fig. 1A). Increased formation of autophagic vesicles was also observed in the polyI:C-transfected and SeV-infected cells compared with that in mock-infected cells, as determined by Cyto-ID that can stain both autophagosomes and autolysosomes specifically (22). The increased autophagosome formation by polyI:C transfection or SeV infection was further confirmed by transmission electron microscopy. As shown in Figure 1C, the number of dense black double-membrane structure vesicles increased in the polyI:C transfected or SeV infected cells. The formation of LC3 puncta by polyI:C or SeV infection also contributed to the modulation of autophagy by RIG-I signaling (Fig. 1D). Furthermore, the level of p62 decreased in a time-dependent manner in the polyI:C-transfected or SeV-infected cells (Fig. 1E). This indicated that the increased level of LC3-II was due to increased autophagy and not due to decreased phagolysosome formation.

**Figure 1.**
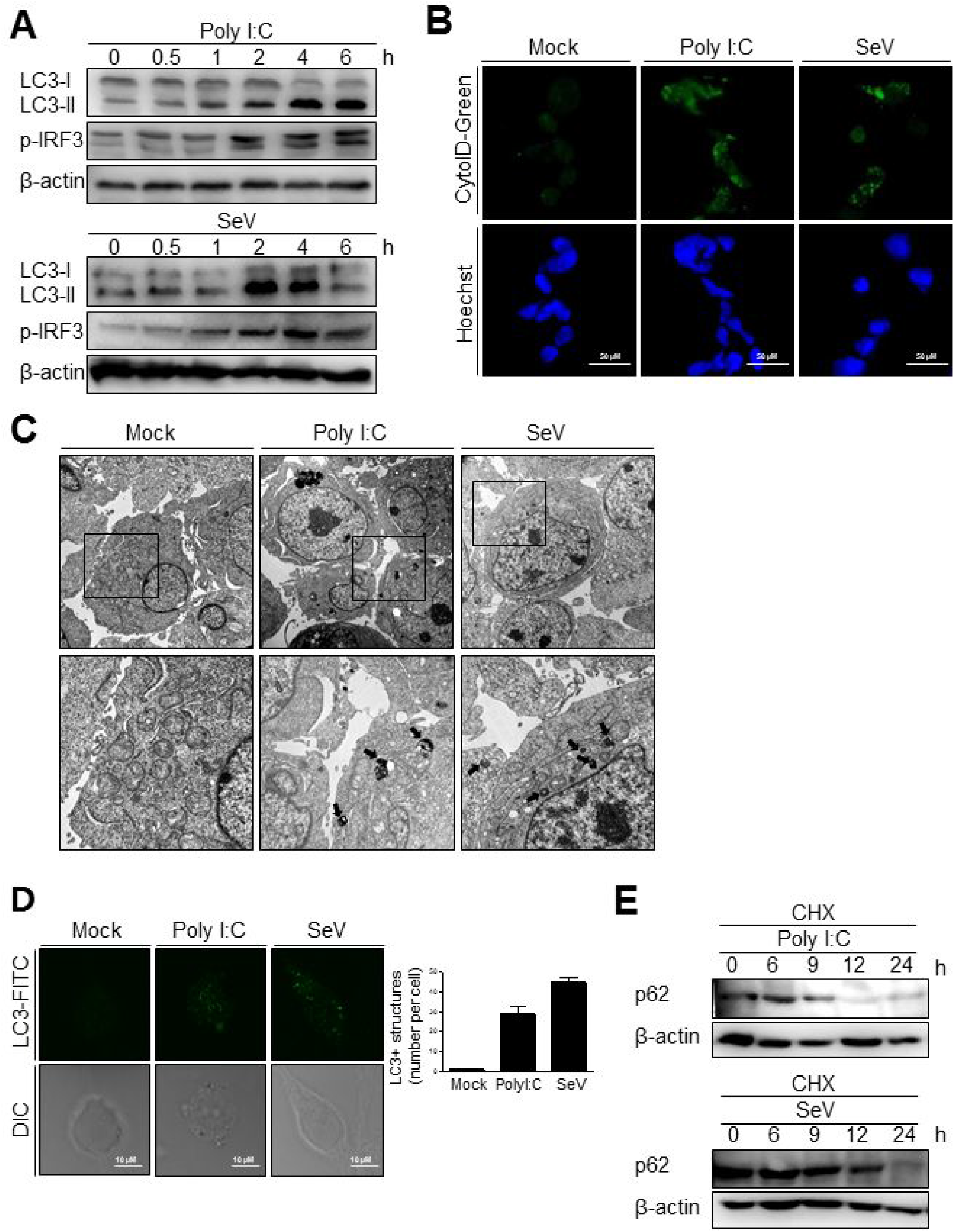
RIG-I activation invokes autophagy. (A) HEK293T cells were transfected with polyI:C (2 μg) or infected with Sendai virus (SeV) (200 HA U/mL) for the indicated hours. The cell lysates were analyzed by immunoblotting using antibodies for LC3, phosphor-IRF3, and β-actin. (B) HEK293T cells were transfected with polyI:C or infected with SeV as in (A) and incubated for 12 h. Autophagosomes were labeled with cytoID-green reagent and observed by fluorescence microscopy. The bottom panels show staining with Hoechst dye to visualize nuclei of the cells. (C) HEK293T cells were transfected with polyI:C (10 μg) or infected with SeV (200 HA U/mL) for 12 h. The cells were harvested, fixed, and subjected to transmission electron microscopy to observe autophagic vesicles (black arrows in the lower panels). The bottom panels show enlarged view of the boxed regions in the top panels. (D) HEK293T cells were mock-treated, transfected with polyI:C or infected with SeV as in (A). After washing and fixation, LC3 puncta were visualized by staining with anti-LC3 antibody and FITC-labeled secondary antibody and observed by fluorescence microscopy. The number of puncta was counted and analyzed using the image J software. The right panel shows the mean number of LC3 puncta in a cell. The data are presented as mean ± standard error of the mean. (E) HEK293T cells were transfected with polyI:C (2 μg) or infected with SeV (200 HA U/mL) and treated with cycloheximide (CHX, 100 ng/mL) for 0, 4, 8, or 12 h. The level of p62 was analyzed by immunoblotting. Each experiment was repeated three or more times.

The induction of autophagy by RIG-I activation was also observed in different types of cells. The transfection of polyI:C was resulted in increased LC3 lipidation and LC3 puncta formation in Raw264.7 murine monocytic cells (Figs. 2A and 2B). Consistently, increment of LC3 lipidation by polyI:C transfection or SeV infection was easily detected in N2a mouse neuroblastoma cells and BV-2 mouse microglial cells (Figs. 2C and 2D), suggesting that the induction of autophagy by RIG-I activation is not a cell- or species-specific phenomenon.

**Figure 2.**
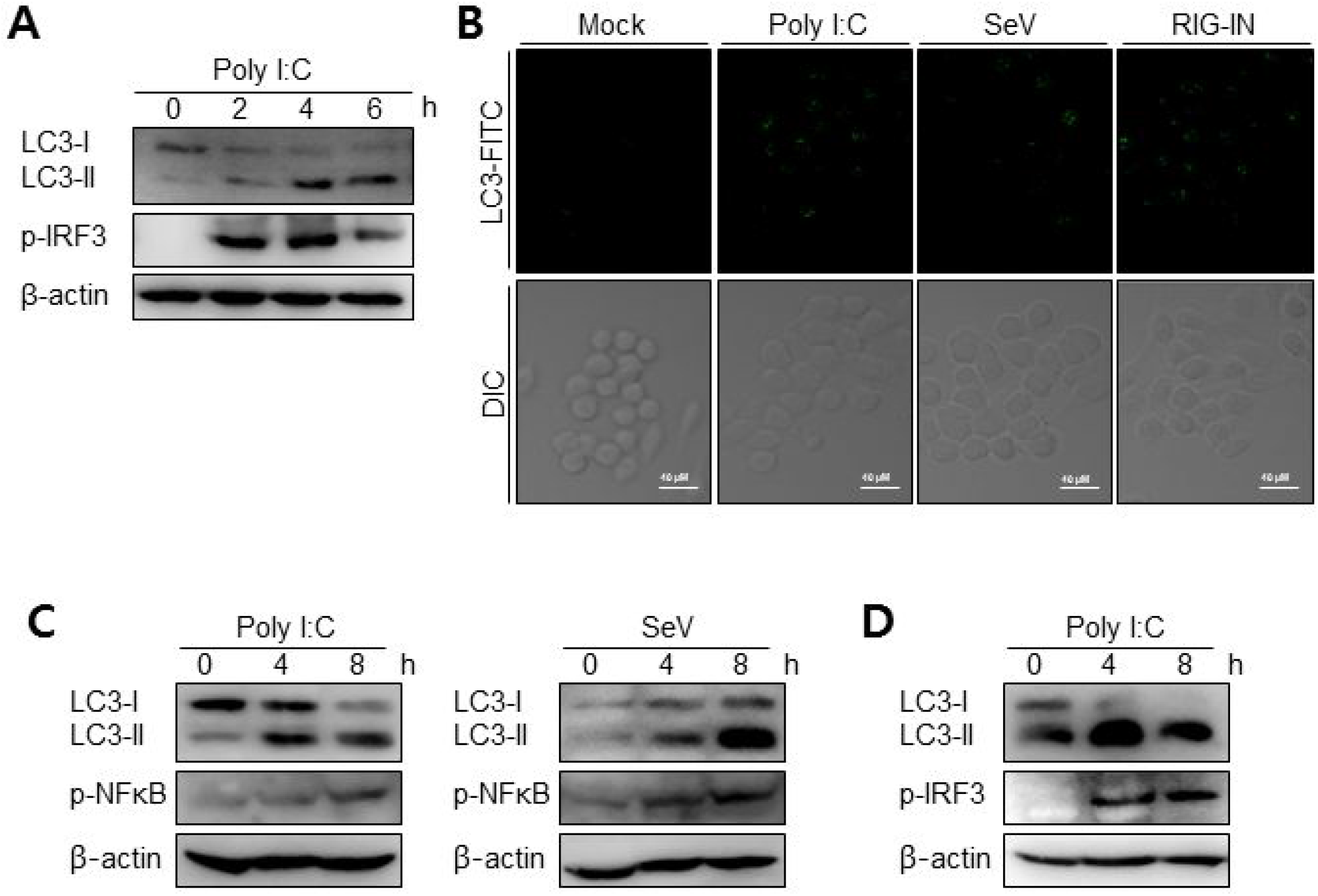
Induction of autophagy by RIG-I in different types of cells. (A) Raw264.7 murine macrophage cells were transfected with polyI:C (2 μg) for the indicated hours. The cell lysates were analyzed by immunoblotting using antibodies for LC3, phospho-IRF3, and β-actin. (B) Raw264.7 cells were mock-treated, transfected with polyI:C (2 μg) or infected with SeV (200 HAU/ml) for 8 h. After washing and fixation, LC3 puncta were visualized by staining with anti-LC3 antibody and FITC-labeled secondary antibody and observed by fluorescence microscopy. (C) N2a murine hypothalamus cells were transfected with polyI:C (2 μg) or infected with Sendai virus (SeV) for 0, 4, or 8 h. The cell lysates were analyzed by immunoblotting using the indicated antibodies as primary antibodies. (D) BV-2 murine microglial cells were transfected with polyI:C and incubated for indicated hours. The cell lysates were analyzed by immunoblotting using indicated antibodies. Each experiment was repeated three or more times.

To further confirm that the triggering of autophagy by an RIG-I agonist was due to the activation of RIG-I signaling, the effect of expressing a constitutively active form of RIG-I on autophagy was examined. The ectopic expression of RIG-I-2Card (RIG-IN) or MDA5-2Card domains (MDA5-N) increased LC3 lipidation (Fig. 3A). The activation of RIG-I signaling by ectopic expression also increased autophagosomes, as determined by cyto-ID staining (Fig. 3B). The ectopic expression of RIG-IN or MDA5-N also increased LC3-functa formation, indicating that both RIG-I and MDA5 can modulate autophagy (Fig. 3C). The level of p62 decreased in the RIG-IN-expressing cells in a time-dependent manner (Fig. 3D). Moreover, the inhibition of lysosomal degradation by the treatment with chloroquine (CQ) significantly increased the effect of RIG-IN on LC3 lipidation, confirming that the accumulation of LC3-II by RIG-I signaling was not due to the suppression of autophagy flux (Fig. 3E).

**Figure 3.**
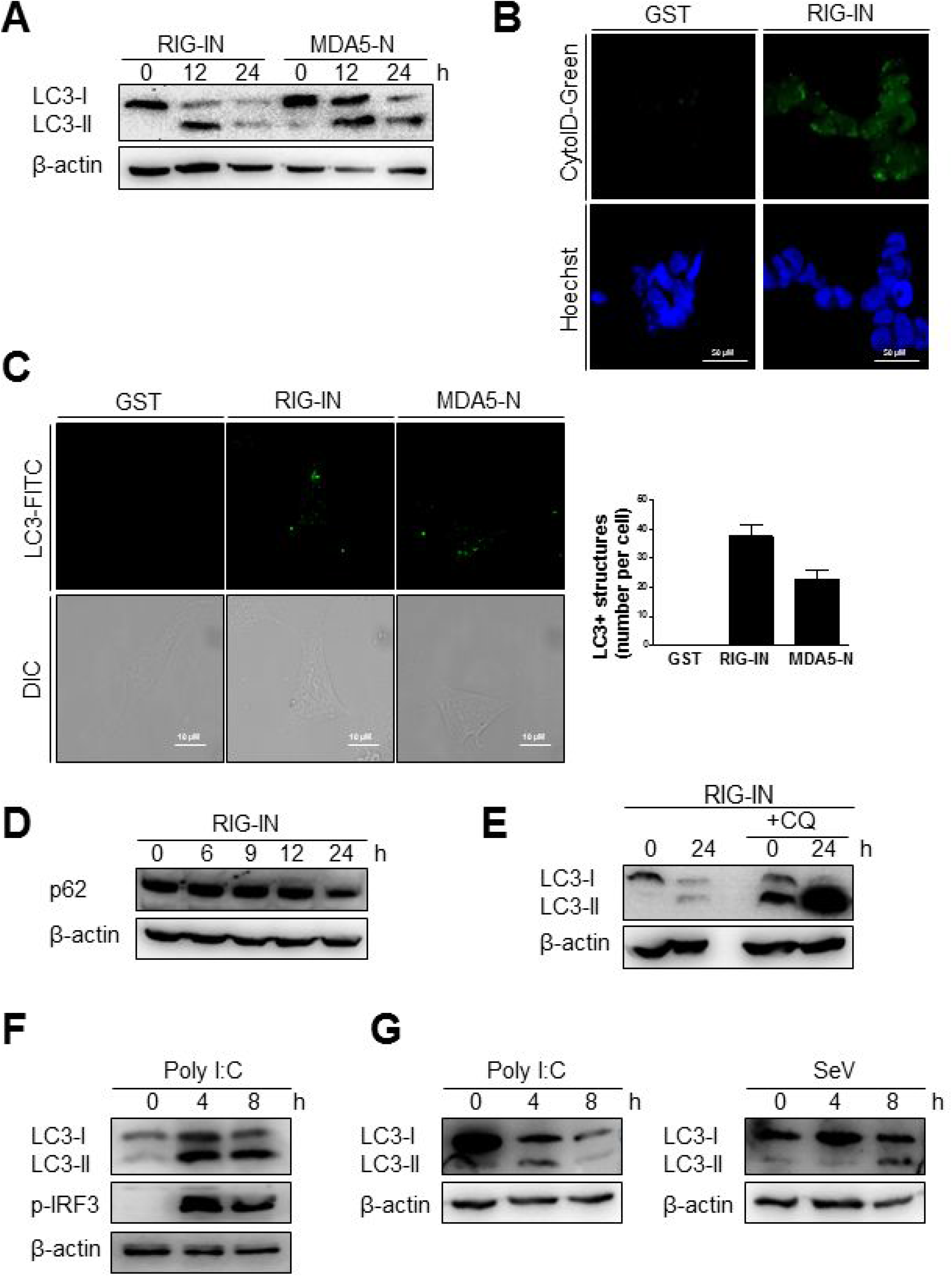
A constitutively active form of RIG-I triggers autophagy. (A) HEK293T cells were transfected with constitutively active N-terminal Card domains of RIG-I (RIG-IN) or MDA5 (MDA5-N). LC3 lipidation was analyzed by immunoblotting using anti-LC3 and anti-β-actin antibodies. (B) HEK293T cells were transfected with vector or RIG-IN and incubated for 24 h. Autophagosomes were stained with Cyto-ID reagent and observed under a fluorescence microscope. The bottom panels show staining with Hoechst dye to visualize nuclei of the cells. (C) HEK293A cells were transfected with RIG-IN or MDA5-N. Eighteen hours after transfection, the cells were fixed and stained with anti-LC3 antibody and FITC-labeled secondary antibody and subjected to fluorescence microscopy. The numbers of puncta was counted and analyzed using the image J software. The right panel shows the mean number of LC3 puncta in a cell. The data are presented as mean ± standard error of the mean. (D) HEK293T cells were transfected with RIG-IN for 0, 6, 9, 12, and 24 h. The level of p62 was analyzed by immunoblotting. (E) HEK293T cells were transfected with vector or RIG-IN for 0 or 24 h with or without chloroquine (CQ) treatment (20 μM) for 12 h before harvest. The cells were subjected to immunoblotting using an anti-LC3 antibody. (F) Type-I interferon receptor (INFR)-deficient mouse embryonic fibroblast cells were transfected with 2 μg polyI:C and incubated for 0, 4, or 8 h. LC3 lipidation was analyzed by immunoblotting. (G) Vero cells were transfected with 2 μg polyI:C or infected with 200 HA U/mL SeV for 0, 4, or 8 h. LC3 lipidation was analyzed by immunoblotting. Each experiment was repeated three or more times.

The induction of autophagy by polyI:C or SeV was examined in cells defective in type-I interferon signaling to test whether RIG-I-mediated autophagy was due to type-I interferon signaling induced by RIG-I signaling. The transfection of polyI:C triggered LC3 lipidation in type-I interferon receptor-deficient MEFs, suggesting that RIG-I signaling can induce autophagy flux in a type-I interferon-independent manner (Fig. 3F). Furthermore, polyI:C transfection or SeV infection also increased LC3-II in Vero cells, which are defective in interferon signaling (Fig. 3G).

### Autophagy induction by SeV infection or polyI:C transfection requires functional RIG-I

The influence of RIG-I-mediated signaling on the induction of autophagy was further examined using cells deficient in RIG-I activity. We used Huh7.5 human hepatoma cells that were derived from Huh7 cells. These cells lose their RIG-I activity due to a mutation in RIG-I (T55I). As expected, LC3-II formation increased by the transfection of Huh7 but not by Huh7.5 cells with polyI:C (Fig. 4A). However, the infection of Huh7.5 cells with influenza A virus activated autophagy to a level comparable to Huh7 cells. This suggested that influenza A virus triggered autophagy via other signaling pathways, such as TLR4, in the absence of RIG-I signaling (Fig. 4B). Consistent with the Huh7.5 data, RIG-I-deficient mouse embryonic fibroblasts (MEFs) also failed to exhibit increased autophagy upon polyI:C transfection or SeV infection, whereas, RIG-I wild-type (WT) MEFs successfully activated autophagy with both treatments (Figs. 4C and D).

**Figure 4.**
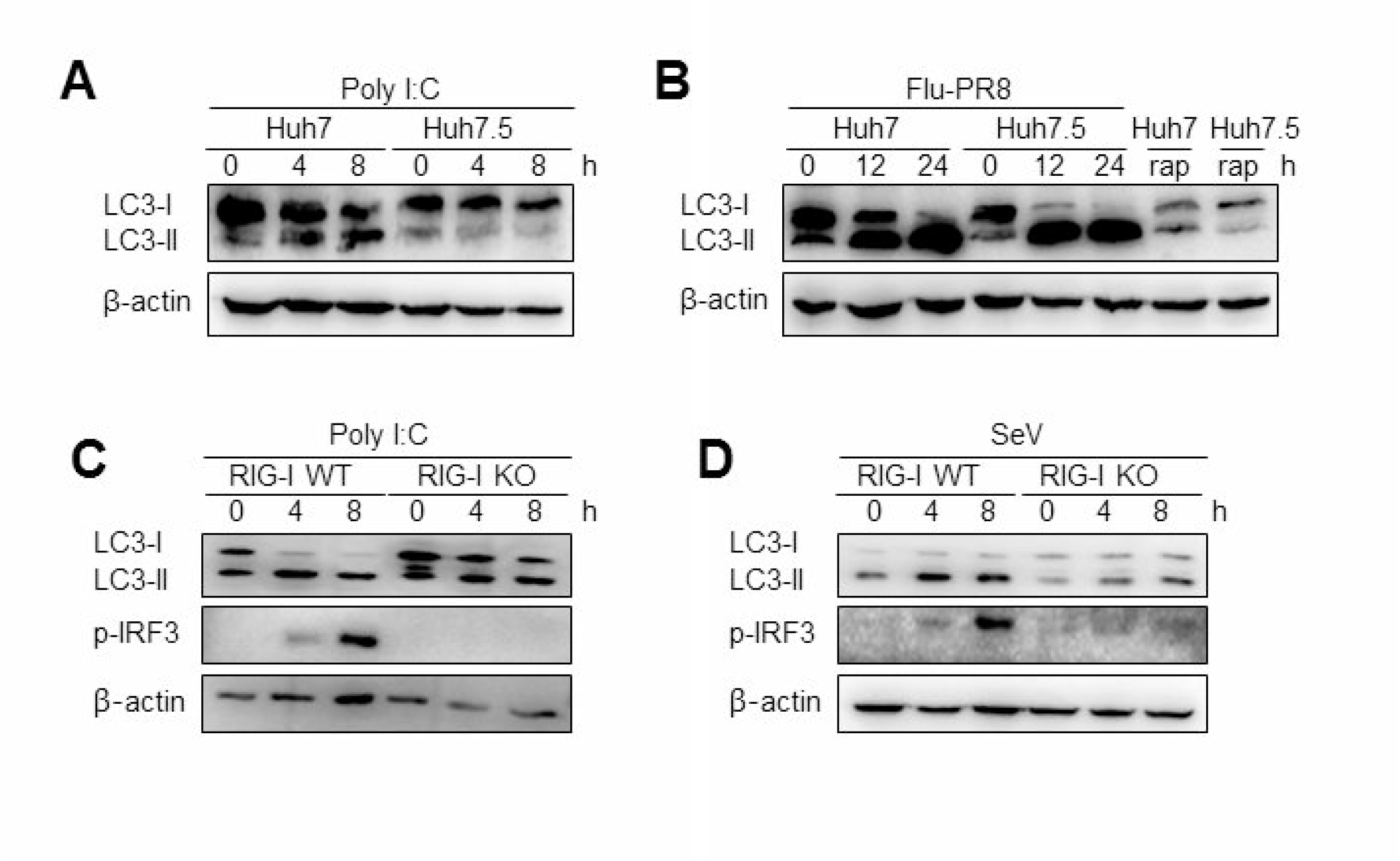
Functional RIG-I is required for polyI:C and Sendai virus (SeV)-mediated autophagy activation. (A) Huh7 human hepatoma cells and Huh7-derived Huh7.5 cells with defective RIG-I activity were transfected with 2 μg polyI:C. LC3 lipidation was analyzed by immunoblotting using anti-LC3 and anti-β-actin antibodies. (B) Huh7 and Huh7.5 cells were infected with influenza A PR8 (Flu-PR8) (multiplicity of infection = 1) as indicated, or were treated with 2 μM rapamycin for 24 h. LC3 lipidation was analyzed as in (A). LC3 lipidation in wild-type (WT) and RIG-I knock-out (KO) mouse embryonic fibroblasts transfected with 2 μg polyI:C (C) or infected with 200 HA U/mL SeV (D). LC3 lipidation was analyzed as in (A). Each experiment was repeated three or more times.

### MAVS-TRAF6 signaling axis is required for RIG-I-mediated signaling

The role of MAVS, a downstream mitochondrial signaling molecule, in inducing autophagy was investigated. The LC3-II level was increased by the ectopic expression of MAVS in HEK293T cells (Fig. 5A). However, the introduction of polyI:C into the MAVS-deficient MEFs failed to increase LC3 lipidation (Fig. 5B). Consistently, no significant increase in autophagosome formation was observed by electron microscopy in the MAVS-deficient MEFs upon transfection with polyI:C. In contrast, a significant increase in the number of autophagosomes was observed in WT MEFs (Fig. 5C). Furthermore, there was no significant decrease in p62 upon SeV infection in the cycloheximide-treated MAVS-deficient MEFs, whereas, a significant decrease in the p62 level was observed in the cycloheximide-treated WT MEFs (Fig. 5D). These results indicate that RIG-I or MDA5 induce autophagy flux via their downstream MAVS. Given that MAVS recruits TRAF6 to activate downstream signaling and TRAF6 activates TLR4-mediated autophagy, it is worth testing the hypothesis that TRAF6 plays a crucial role in RIG-I-mediated autophagy. Indeed, LC3-II formation following SeV infection or polyI:C transfection was significantly lower in TRAF6-deficient than in WT MEFs (Figs. 6A and B). Consistently, LC3 puncta formation by transfection with polyI:C or RIG-IN was significantly lower in the TRAF6-defective MEFs than in the WT MEFs (Fig. 6C). In addition, there was a significant decrease in the level of p62 in the SeV-infected WT cells but not in the TRAF6 knock-out (KO) MEFs (Fig. 6D).

**Figure 5.**
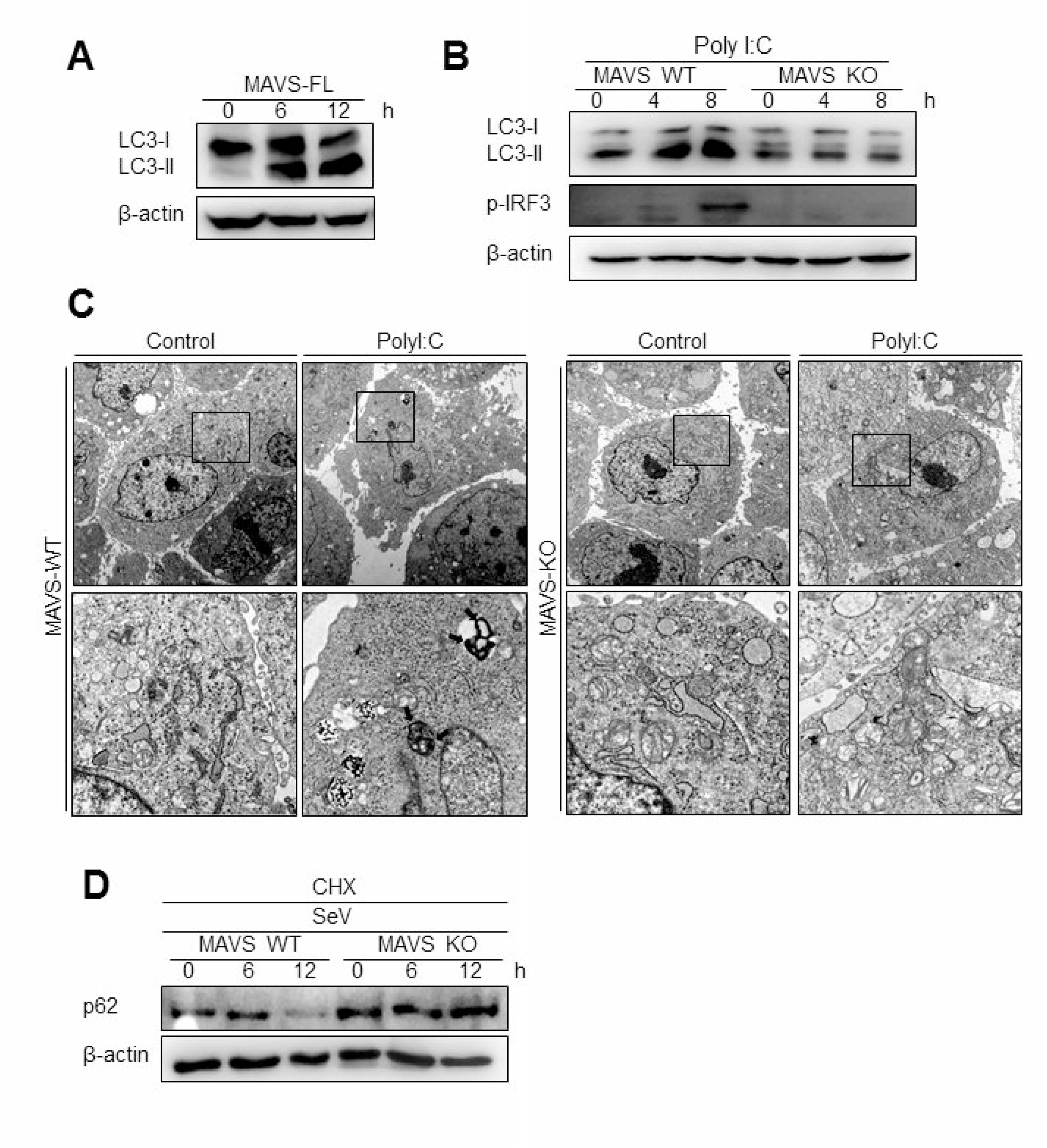
Mitochondrial antiviral signaling protein (MAVS) is required for the induction of autophagy. (A) HEK293T cells were transfected with MAVS and harvested after 0, 6, and 12 h. LC3 lipidation was analyzed by immunoblotting using an anti-LC3 antibody. (B) The wild-type (WT) and MAVS knockout (KO) mouse embryonic fibroblasts (MEFs) were transfected with 2 μg polyI:C for 0, 4, or 8 h. LC3 lipidation was analyzed by immunoblotting as in (A). (C) The WT and MAVS KO MEFs were transfected with 2 μg polyI:C for 12 h. The cells were observed by transmission electron microscopy. The bottom panels show enlarged view of the boxed regions in the top panels. (D) The WT and MAVS KO MEFs were infected with 200 HA U/mL SeV and treated with cycloheximide (CHX, 100 ng/mL) for 0, 6, or 12 h. The level of p62 was analyzed by immunoblotting. Each experiment was repeated three or more times.

**Figure 6.**
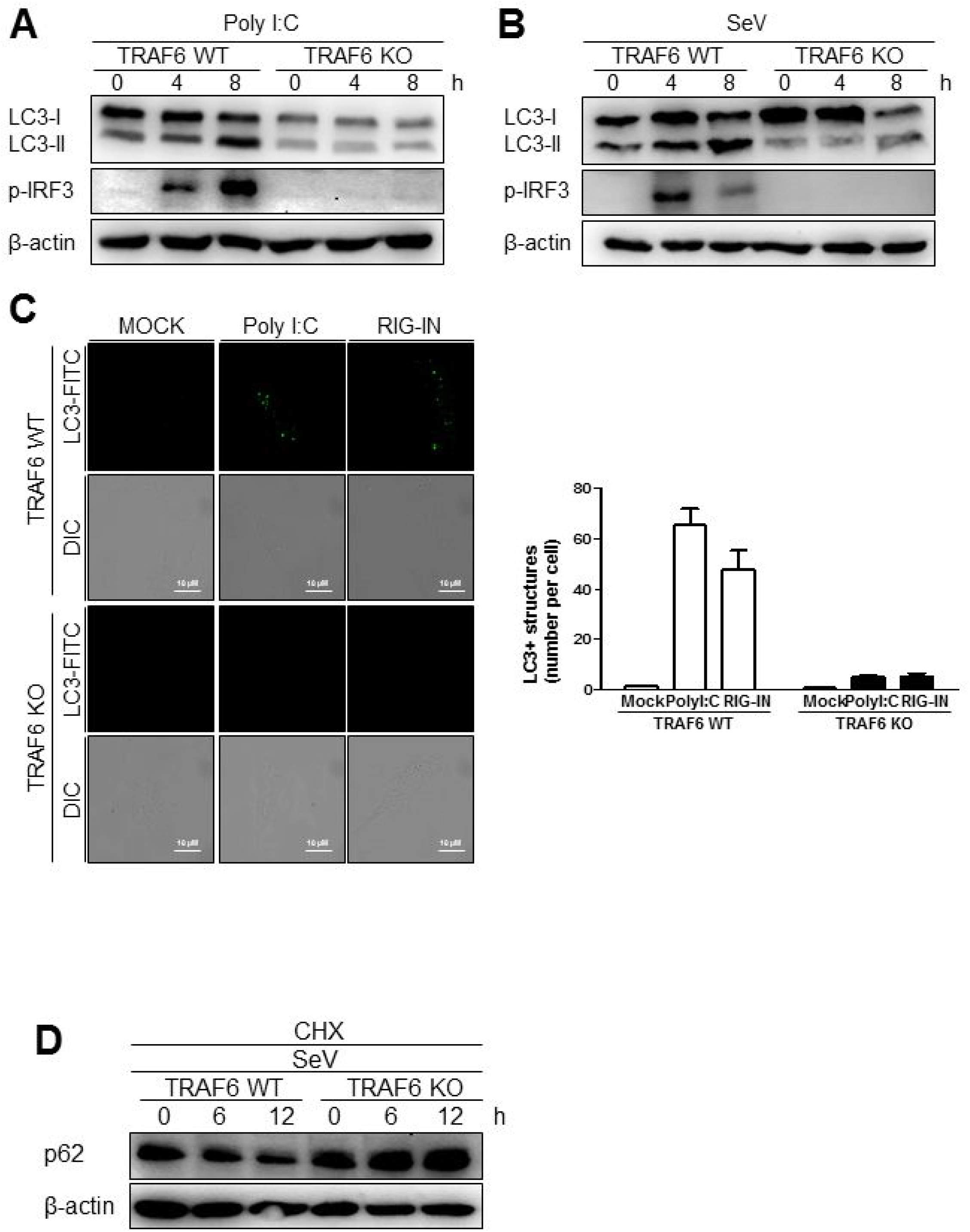
Tumor necrosis factor receptor-associated factor (TRAF)6 is required for the induction of autophagy. The WT and TRAF6 KO MEFs were transfected with 2 μg polyI:C (A) or infected with 200 HA U/mL SeV (B) for 0, 4, or 8 h. LC3 lipidation was analyzed by immunoblotting using an anti-LC3 antibody. (C) The TRAF6 WT and KO MEFs were transfected with polyI:C (2 μg/well) or RIG-IN (100 ng/well). Eighteen hours after transfection, LC3-puncta was visualized by staining with anti-LC3 antibody and FITC-labeled secondary antibody. The number of puncta was counted and analyzed using the image J software. The right panel shows the mean number of LC3 puncta in a cell. Data presented as mean ± standard error of the mean. (D) The WT and TRAF6 KO MEFs were infected with 200 HA U/mL SeV and treated with CHX (100 ng/mL) for 0, 6, or 12 h, then subjected to immunoblotting using an anti-p62 antibody. Each experiment was repeated three or more times.

### TRAF6 associates with Beclin-1 upon RIG-I activation

TRAF6 interacts with Beclin-1 to activate TLR4-mediated autophagy. Thus, we explored the possible interaction between TRAF6 and Beclin-1 upon RIG-I activation. The interaction between overexpressed Beclin-1 and endogenous TRAF6 was detected by the co-immunoprecipitation (co-IP) assay. The interaction increased significantly by the ectopic expression of RIG-IN (Fig. 7A). The transfection of polyI:C into V5-Beclin-1-expressing cells, followed by co-IP experiment showed an increase in the interaction of Beclin-1 with the autophagy initiation complex components, including VPS34, ATG14, and Ambra-1, indicating that Beclin-1 autophagy initiation complex formation is triggered by RIG-I activation (Fig. 7B). Furthermore, the co-IP assay showed an association between endogenous Beclin-1 and TRAF6 upon transfection of polyI:C. This assay also demonstrated that VPS34 interacted with Beclin-1 upon polyI:C transfection (Fig. 7C). These results suggest that Beclin-1 associates with VPS34 and TRAF6 to facilitate autophagy upon RIG-I activation. Thus, we examined whether Beclin-1 migrated to mitochondria upon RIG-I activation to interact with TRAF6, which can recruit mitochondrial MAVS. As shown in Figure 7D, the translocation of Beclin-1 to mitochondria was detected in the cells overexpressing RIG-IN, indicating that Beclin-1 can be recruited to mitochondria upon activation of RIG-I signaling. Increased localization of Beclin-1 in the mitochondria upon RIG-IN expression or SeV infection was also confirmed by co-localization of Beclin-1 with mitotracker using confocal microscopy (Figs. 7E and 7F).

**Figure 7.**
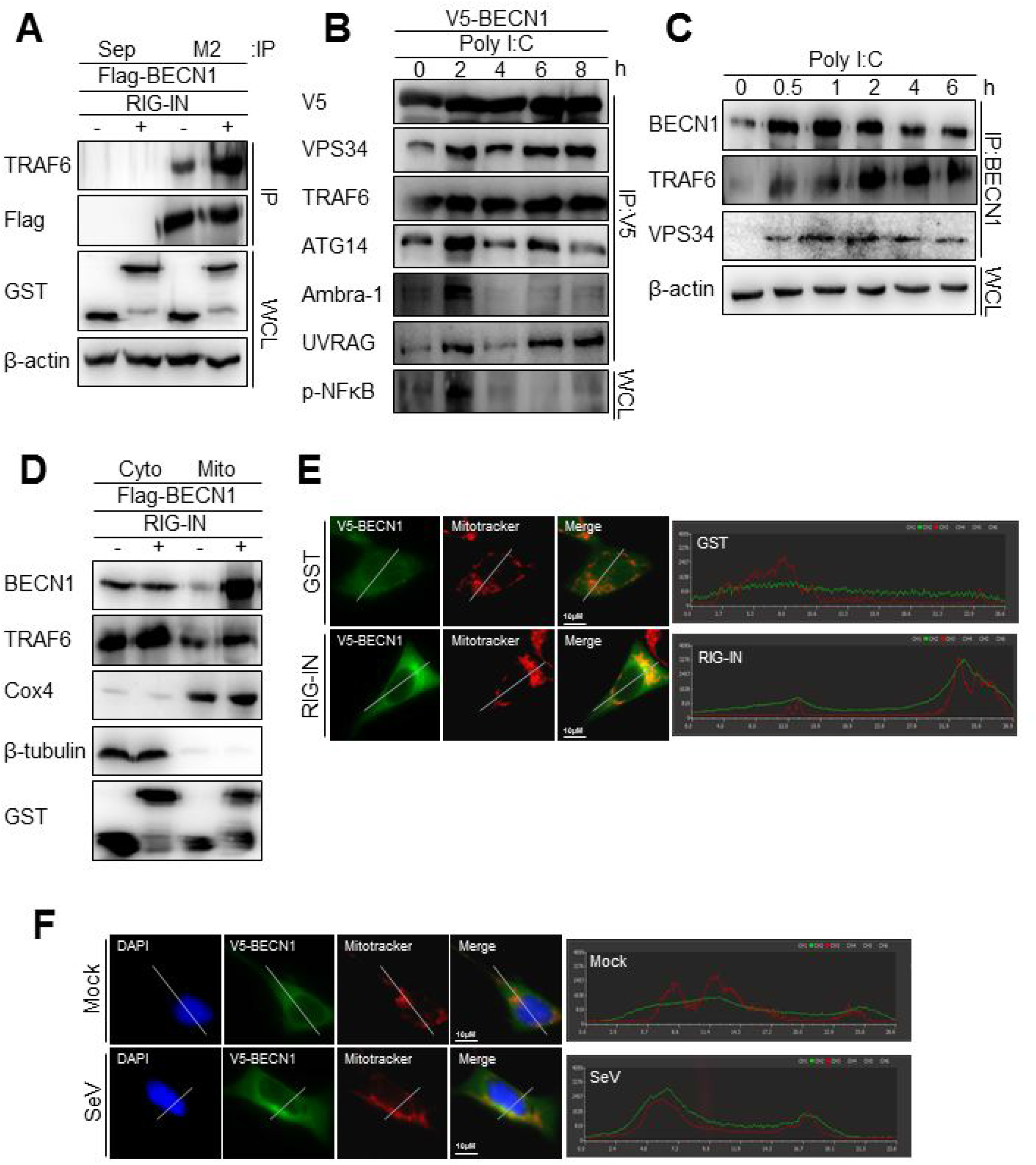
RIG-I activation increases the interaction between tumor necrosis factor receptor-associated factor (TRAF)6 and Beclin-1. (A) HEK293T cells were transfected with Flag-Beclin-1 (Flag-BECN1) and pEBG (GST) or pEBG-RIG-IN. Thirty-six hours after transfection, the cell lysates were subjected to co-immunoprecipitation (co-IP) using a M2 anti-Flag antibody-coated resin. The sepharose resin (Sep) served as a negative control. Whole cell lysates (WCL) and samples from co-IP were analyzed by immunoblotting using the indicated antibodies. (B) HEK293T cells were transfected with V5-Beclin-1. Twelve hours after transfection, polyI:C was transfected into the cells at different time points and incubated for the indicated hours. The cell lysates were subjected to co-IP using anti-V5 antibody, followed by immunoblotting using the indicated antibodies. (C) HEK293T cells were transfected with polyI:C and harvested at 0, 0.5, 1, 2, 4, and 6 h. The cell lysates were subjected to co-IP with an anti-Beclin-1 (BECN1) antibody and analyzed by immunoblotting using the indicated antibodies. (D) HEK293T cells were transfected with Flag-BECN1, pEBG (GST), or pEBG-RIG-IN and incubated for 24 h. The mitochondrial and cytoplasmic fractions were separated as described in the Materials and Methods and subjected to immunoblotting using the indicated antibodies. (E) HEK293A cells were transfected with GST control vector or GST-RIG-IN together with V5-Beclin-1. Eighteen hours after transfection, the cells were stained with Mitotracker and anti-V5 antibody as described in the Materials and Methods. Fixed dishes were observed by confocal microscopy. The right panel shows the intensity of Mirotracker (Red) and Beclin-1 (Green) along the white line of the left panel images. (F) HEK 293T cells were transfected with V5-Beclin-1. Sixteen hours after transfection, the cells were mock-infected or infected with SeV for 2h. The localization of Beclin-1 was analyzed by confocal microscopy as in (E). Each experiment was repeated three or more times.

### Beclin-1 undergoes K63-linked polyubiquitination upon RIG-I activation

As TRAF6 is an E3-ubiquitin ligase, and K63-linked polyubiquitination of Beclin-1 modulates its function in autophagy, we hypothesized that the interaction between Beclin-1 and TRAF6 might lead to K63-linked polyubiquitination of Beclin-1. Indeed, the ubiquitination of TRAF6 was increased by the ectopic expression of RIG-IN and further increased by the treatment with CQ, as determined by IP of the overexpressed TRAF6 and immunoblotting using an anti-ubiquitin antibody (Figs. 8A and 8B). Immunoblotting using the K63-linked ubiquitin-specific antibody showed that the K63-ubiquitination of TRAF6 and its interaction with VPS34 were also increased by RIG-IN expression and CQ treatment (Fig. 8C). PolyI:C transfection also increased the ubiquitination of Beclin-1 within 30 min (Fig. 8D).

**Figure 8.**
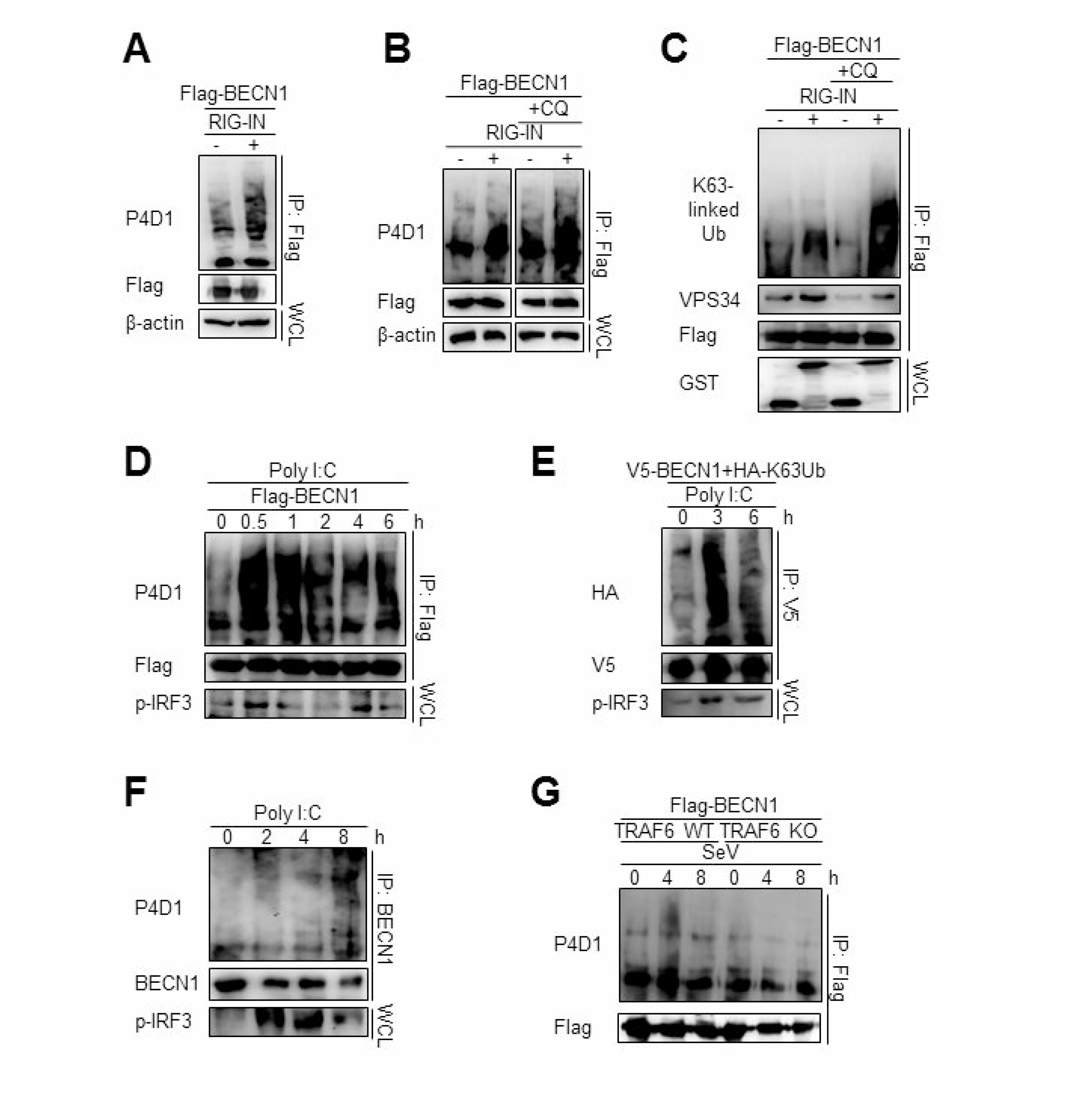
RIG-I activation increases K63-linked polyubiquitination of Beclin-1. (A) HEK293T cells were transfected with Flag-Beclin-1 (Flag-BECN1) and pEBG (GST) or pEBG-RIG-IN, and incubated for 24 h. The cell lysates were subjected to immunoprecipitation (IP) using M2 anti-Flag resin and analyzed by immunoblotting using the indicated antibodies. (B) HEK293T cells were transfected with Flag-BECN1 and pEBG-RIG-IN, and treated with or without 20 μM chloroquine (CQ). The cell lysates were analyzed by IP and immunoblotting as in (A). (C) HEK293T cells were transfected with the indicated plasmids and incubated in the presence or absence of 20 μM CQ for 12 h. Lys63 (K63)-linked polyubiquitination of Beclin-1 was analyzed by IP and immunoblotting with the indicated antibodies. (D) HEK293T cells were transfected with Flag-Beclin-1. After 24 h, the cells were transfected with polyI:C and incubated for 0, 0.5, 1, 2, 4, and 6 h. The cell lysates were analyzed by IP and immunoblotting. (E) HEK293T cells were transfected with Flag-Beclin-1 and HA-K63-only ubiquitin mutant plasmids. After 24 h, the cells were transfected with polyI:C and incubated for 0, 3, or 6 h. The cell lysates were analyzed by IP and immunoblotting. (F) HEK293T cells were transfected with polyI:C and incubated for 0, 2, 4, and 8 h. The cell lysates were subjected to IP using the anti-Beclin-1 (BECN1) antibody and analyzed by immunoblotting. (G) Wild-type (WT) and TRAF6 knock-out (TRAF6 KO) MEFs were transfected with Flag-BECN1. After 24 h, the cells were infected with SeV and incubated for 0, 4, or 8 h. The cell lysates were subjected to IP and immunoblotting. Each experiment was repeated three or more times.

To further confirm the K63-linked polyubiquitination of Beclin-1 upon RIG-I activation, we used a K63-only ubiquitin mutant where all lysines, except K63, were mutated to arginine and thereby restricting the polyubiquitination to K63. The PolyI:C transfection of HEK293T cells ectopically expressing the HA-tagged K63-only ubiquitin mutant with Beclin-1 showed increased K63-linked polyubiquitination of Beclin-1 (Fig. 8E). In addition, the ubiquitination of endogenous Beclin-1 upon polyI:C transfection was observed by IP and immunoblotting (Fig. 8F). To determine whether TRAF6 was required for RIG-I-mediated Beclin-1 ubiquitination, the WT and TRAF6 KO MEFs were analyzed. As shown in Figure 8G, there was no significant change in the Beclin-1 ubiquitination level in the TRAF6 KO MEFs upon SeV infection, whereas, increased ubiquitination was detected in the WT MEFs. Collectively, these results show that Beclin-1 undergoes K63-linked polyubiquitination upon RIG-I activation in a TRAF6-dependent manner to facilitate autophagy flux.

## Discussion

In addition to its basal roles in maintaining cellular homeostasis, autophagy has been implicated in immunity. Autophagy affects the (1) innate and adaptive immune systems by directly eliminating pathogens, controlling inflammation, and facilitating antigen presentation and (2) secretion of immune mediators (23). Autophagy is triggered by infection with a diverse range of viruses suggesting there is crosstalk between the innate immune recognition of viral infection and the autophagy pathway (23). Recent studies have revealed that the innate immune sensors, including TLRs and cGAS, can trigger the autophagic process (16, 17). Regarding RLRs, a deficiency of autophagy augments RIG-I-mediated type-I interferon and some autophagic proteins, such as the ATG5-ATG12 complex, that inhibit the RIG-I signaling pathway. This suggests a negative regulatory role of autophagy in the RIG-I-mediated signaling pathway (21–24). Recent studies have showed that polyI:C transfection activates autophagy and MAVS maintains mitochondrial homeostasis via autophagy (25). However, it is not clear whether RIG-I activation by the recognition of a pathogen-associated molecular pattern regulates the autophagic process directly.

The results of the present study provide evidence that RIG-I triggers autophagic flux upon recognition of its ligands. The induction of autophagy following the intracellular introduction of a synthetic RNA (polyI:C), SeV infection, or forced expression of a constitutively active form of RIG-I, clearly indicates the importance of RIG-I signaling. Moreover, the RIG-I ligands failed to induce autophagy in cells defective in RIG-I signaling. The main effect of RIG-I-mediated antiviral signaling is the production of type-I interferon, which can facilitate autophagy (26, 27). Surprisingly, activating RIG-I increased autophagy in type-I interferon signaling-defective cells, indicating that RIG-I signaling-mediated autophagy is independent from type-I interferon-mediated autophagy.

A crucial role of TRAF6 during autophagy induced by the innate immune system has been shown by several studies. In macrophages, activated CD40 recruits TRAF6 and induces Beclin-1-dependent autophagy to eliminate infected *Toxoplasma gondii* (28, 29). In addition, TLR4 signaling requires TRAF6-mediated Beclin-1 ubiquitination to induce autophagy. This suggests that the recruitment and activation of TRAF6 by innate immune signaling may lead to the ubiquitination of Beclin-1 and formation of downstream signaling complexes, including VPS34 (17).

Our study using MAVS- and TRAF6-deficient cells has proven the crucial role of the MAVS-TRAF6 signaling axis in the RIG-I-dependent pathway. We demonstrated that TRAF6 is required for RIG-I-mediated autophagy, and Beclin-1 is translocated to the mitochondria, and it interacts with TRAF6 upon RIG-I activation. These data suggest that RIG-I may share TRAF6-dependent downstream signaling with TLR4 signaling to promote autophagy. Notably, Beclin-1 interacts simultaneously with VPS34 and TRAF6 upon polyI:C transfection, suggesting the possible role of TRAF6 in the formation of an active Beclin-1 complex.

Several recent studies have showed that the K63-linked polyubiquitination of Beclin-1 is crucial for autophagy activation and can be targeted by cellular proteins to modulate autophagy (17, 30, 31). Our data suggest that the TRAF6-mediated K63-polyubiquitination of Beclin-1 upon RIG-I activation may activate the Beclin1-VPS34 complex to induce autophagy. In a previous study, Beclin-1 was shown to interact with MAVS to suppress RIG-I-mediated interferon production in an ATG-5-independent manner (32). Our data showing the mitochondrial translocation of Beclin-1 upon RIG-I activation suggest that this may have the dual effect of suppressing the RIG-I interaction with MAVS and inducing autophagy to negatively regulate RIG-I signaling.

Collectively, RIG-I activation leads to Beclin-1 K63-polyubiquitination and mitochondrial translocation to induce autophagy in a MAVS-TRAF6-dependent manner. Given that autophagy can suppress RIG-I-mediated interferon production, it seems likely that RIG-I-induced autophagy serves as a negative-feedback mechanism to prevent an excessive response. It would be interesting to examine whether viral components, such as nucleic acids, proteins, or viral particles in the cytoplasm, can be targeted for the autophagic elimination process upon RIG-I activation.

## Materials and Methods

### Cells, viruses, and plasmids

The RIG-I KO and WT MEFs were described previously (33). The MAVS KO and IFNAR KO MEFs were generated from the MAVS KO mice (kindly provided by Dr. Shizuo Akira at the Osaka University) and IFNAR KO mice (kindly provided by Dr. Heung Kyu Lee at the Korea Advanced Institute of Science and Technology). The TRAF6 KO MEFs were kindly provided by Dr. Yoon-Jae Song at the Gachon University. N2a and BV-2 cells were described preciously (34). HEK293T, HEK293A, Raw264.7, Huh7, Huh7.5, Vero and MEF cells were maintained in Dulbecco’s modified Eagle’s medium containing 10% fetal bovine serum and penicillin/streptomycin (100 U/mL). SeV (Cantell strain) was purchased from Charles River Laboratories. Influenza A virus was prepared and infected as described previously (35).

### Immunoblotting and co-IP

The cells were transfected with the indicated plasmids and treated as described. The cells were collected and lysed with Triton X-100 lysis buffer (25 mM Tris-HCl, pH 7.5, 150 mM NaCl, 1 mM EDTA, 0.5% Triton X-100) containing protease inhibitor cocktail (Pierce, #78430). After centrifugation, the cell lysates were subjected to sodium dodecyl sulfate polyacrylamide gel electrophoresis (SDS-PAGE), followed by immunoblotting. Primary antibodies used were as follows: anti-LC3 (Cosmo Bio, #CTB-LC3-1-50), anti-SQSTM1/P62 (Abcam, #ab56416, Cambridge, UK), anti-phospho-IRF3 (Cell Signaling Technology, #4947), anti-phospho-NFκB (Cell Signaling Technology, #3031), anti-β-actin (Santa Cruz Biotechnology, #sc-47778, Dallas, TX, USA), anti-ubiquitin P4D1 (Cell Signaling Technology, #3936), anti-Flag (Sigma-Aldrich, #F7435, Saint Louis, MO, USA), anti-Beclin-1 (Cell Signaling Technology, #3738), anti-HA (Santa Cruz Biotechnology, #sc-7392), anti-V5 (Cell Signaling Technology, #13202), anti-K63-linked ubiquitin (Cell Signaling Technology, #5621), anti-VPS34 (Cell Signaling Technology, #4263), anti-ATG14 (Cell Signaling Technology, #96752), anti-Ambra-1 (Cell Signaling Technology, #12250), anti-UVRAG (Cell Signaling Technology, #5320), anti-GST (Abcam, #ab19256, Cambridge, UK), anti-TRAF6 (Cell Signaling Technology, #8028), anti-Cox4 (Santa Cruz Biotechnology, #133478), anti-β-tubulin (Cell Signaling Technology, #2146). For IP, the clarified cell lysates were incubated with the indicated antibodies for 12 h at 4°C, followed by further incubation with protein A/G resin for 2 h. For IP Flag-tagged proteins, the cell lysates were incubated with anti-Flag M2 affinity resin (Sigma-Aldrich, #A2220) or anti-V5 affinity resin (Sigma-Aldrich, #A7345) for 12 h at 4°C. After extensive washing with lysis buffer, the bound proteins were suspended in 1× sample buffer and analyzed by SDS-PAGE and immunoblotting.

### Autophagosome staining by Cyto-ID and LC3 antibodies

HEK293T cells were transfected with 2 μg polyI:C or infected with SeV at 100 HA units/mL, and collected after 12 h. Subsequently, Cyto-ID autophagy reagent (Enzo, #ENZ-51031, Farmingdale, NY, USA) staining was performed according to the instruction of the manufacturer. Briefly, the cells were washed twice with 1× assay buffer and treated with diluted Cyto-ID green stain solution. The cells were incubated for 30 min at 37°C, and then washed and incubated for 20 min with 4% paraformaldehyde. For LC3 staining, the cells were fixed with 4% paraformaldehyde and permeabilized using 0.1% tritonX-100 buffer. Then cells were stained with LC3 antibody (Cell Signaling Technology, #3868) for 2 h at 37°C and stained with anti-rabbit IgG FITC reagent. The cells were then washed three times and analyzed by fluorescence microscopy to observe punctated forms of autophagosomes.

### Transmission electron microscopy

HEK293T cells were transfected with 10 μg polyI:C using Lipofectamine 2000, or infected with SeV at 100 HA units/mL, and incubated for 12 h. The WT and MAVS KO MEF cells were transfected with polyI:C (10 μg), pEBG vector (2.5 μg), or pEBG-RIG-IN (5 μg) using Lipofectamine 2000, and further incubated for 12 h. The cells were fixed with 2% paraformaldehyde and 2.5% glutaraldehyde in 0.1 M phosphate buffer (pH 7.2) at 4°C for 24 h. The cells were then embedded in epoxy resin and polymerized at 38°C for 12 h, followed by further incubation at 60°C for 48 h. Thin sections, cut using an ultramicrotome (MT-XL, RMC Products), were collected on a copper grid and stained with 4% lead citrate and saturated 4% uranyl acetate. The samples were examined at 80 kV with a transmission electron microscope (JEM-1400Plus, JEOL, Tokyo, Japan) at the Seoul National University Hospital Medical Research Institute (Seoul, Korea). Double- or single-membrane vesicles measuring 0.3 to 2.0 μm in diameter were defined as autophagosomes.

### Mitochondria isolation

HEK293T cells (1 × 10^7^) were transfected with the pEBG vector or pEBG-RIG-IN and incubated for 24 h. The cells were washed with phosphate-buffered saline and mitochondria were isolated using a kit (Thermo Scientific, #89874) according to the instruction of the manufacturer instructions. The cytosolic and mitochondrial fractions were analyzed by SDS-PAGE and immunoblotting. Cox4 and β-tubulin served as markers for the cytosolic and mitochondrial fractions, respectively.

### Immunofluorescence staining and confocal microscopy

HEK293A cells were transfected with polyI:C or treated with SeV as described. The cells were moved to fibronectin-coated confocal dish and incubated for 12 h. The cells were stained with mitotracker (Thermo Scientific, #M7512, Rockford, IL, USA) according to the instruction of the manufacturer and fixed with 4% paraformaldehyde for 15 min, permeabilized using 0.1% tritonX-100 for 10 min and blocked with 5% BSA. The cells were then stained with the primary and secondary antibodies (Thermo Scientific, #31556), (Thermo Scientific, #62-6511) according to the instruction of the manufacturer. The colocalization images were examined under an Olympus FV-1000 confocal microscope.

### Statistical analysis

Data are represented as mean ± standard error of the mean (SEM) unless otherwise indicated, and were analyzed by Student’s unpaired two-tailed *t* test using GraphPad Prism 5 software. A value of p < 0.05 was considered significant.

## Acknowledgments

This work was supported by the Basic Science Research Program grants through the National Research Foundation of Korea (NRF), which was funded by the Ministry of Science, ICT and Future Planning (NRF-2017R1A2B4005596). This work was also supported by the Medical Research Center Program through the National Research Foundation of Korea funded by the Ministry of Science and ICT (NRF-2017R1A5A201476).

